# Mitigative effects of Alpha-lipoic acid for the toxicity of Dimethoate in male rats

**DOI:** 10.1101/527283

**Authors:** Hani M. Abdelsalam

## Abstract

**Background:** Organophosphates are widely used by human nowadays, but these compounds have tremendous negative effect on the man health. So this study aims to use of Alpha-lipoic acid (ALA) to alleviates the negative effects of Dimethoate (DM).

**Methods:** This study is designed as follows, Thirty adult male Wistar albino rats were utilized, further subdivided into control, DM and DM+ALA groups. Liver and renal cortex sections from all groups were processed for histopathological examination, biochemical estimation of liver function tests, serum Urea, Creatinine, BUN, testosterone and lipid profile were performed.

**Results:** This study clarified the improvement effects of ALA against the negative effects of DM where ALA caused a significant recovery of the hepatic (ALT, AST, ALP and total protein) and renal functions by normalizing them in DM + ALA group and to some extent improvement of lipid profile and testosterone levels. Also, ALA restored normal hepatic and renal histomorphology.

**Conclusion:** It is concluded that ALA therapy can ameliorate the negative effects of DM that affect the vital organs as the liver and kidney. Also ALA can reduce the occurrence of atherogensis by reducing the levels of bad cholesterol in the blood. ALA boosts the levels of testosterone so it augments the male sexual characters.

## 1. Introduction

The organophosphates (OPs) like dimethoate (DM) are a serious threat since World War II and have major contribution in an estimated thirty thousand severe pesticide poisoning reported worldwide [1]. Another major implication of this group is the inhibitory effect on acetylcholinesterase -AChE, [2], and its possible implication as it induces changes in characteristics of oxidative stress [3]. These compounds in general, both in chronic and acute intoxication disturb the redox processes, thereby changing the activities of antioxidative enzymes processes and causing enhancement of lipid peroxidation in many organs [4]. The same appeared to be the case for dimethoate, since these compounds are known to increase the production of reactive oxygen species – ROS and attenuation of the antioxidant barrier of the organism so as to induce oxidative stress [1].

Repeated or prolonged exposure to organophosphates may result in many effects as acute exposure, including the delayed symptoms. Workers repeatedly exposed to DM reported impaired memory, disorientation, depressions, irritability, confusion, headache, speech difficulties, delayed reaction times, nightmares, sleepwalking and drowsiness or insomnia. Also, influenza-like condition with headache, nausea, weakness, loss of appetite, and malaise has also been reported [5]. Dimethoate affects the functions of multiple organs including liver. Dimethoate was reported to alter the level of the marker parameters related to the liver in rats and mice [6]. The liver is the primary organ involved in metabolism and is a target organ of OPs and drugs. Clinical biochemistry and hispathological evaluations of liver are the used methods for detecting OP exposure effect [7].

Alpha-lipoic acid (ALA) is a naturally occurring dithiol compound that functions as an essential cofactor for mitochondrial bioenergetic enzymes [8]. It has also been recognized as a powerful antioxidant capable of prevention or treatment of many diseases associated with oxidative stress, such as diabetes, chronic liver diseases, and neurodegenerative processes [9]. ALA supplementation attenuates renal injury in rats with obstructive nephropathy and further suggest that oxidative stress inhibition is likely to be involved in the beneficial effects of this compound.[10]. The therapeutic potential of ALA has been demonstrated in a variety of disorders linked to oxidative stress and inflammation in diverse organ systems, including the kidney [11].

ALA and its reduced dithiol form, dihydrolipoic acid (DHLA), are powerful antioxidants that can scavenge hydroxyl radicals, singlet oxygen, hydrogen peroxide, hypochlorous acid, peroxynitrite, and nitric oxide, and the category is much broader than those of Vit E, ALA/DHLA redox couple can regenerate exogenous and endogenous antioxidants such as Vit C and E, and GSH.Also, the couple exerts additional antioxidant actions through the chelation of copper, iron, and other transitional metals [12]. ALA giving caused a retraction of histopathologic and morphologic lesions in extracted rat kidney. Also, positive effect of ALA on albuminuria was observed. However, no positive effect of ALA was found for antioxidant parameters at the tissue level [13].

## 2. Material and Methods

### 2.1 Experimental animals

Thirty healthy adult male Wistar albino rats (11–13 weeks) weighing 160–180 g were obtained and maintained at the Breeding Animal House of the Faculty of Medicine, Zagazig University, Egypt. Animals were kept for acclimatization in plastic cages with stainless steel wire bar lid at a controlled temperature (23±1°C) and humidity (55±5%) in a (12:12 h light: dark cycle) artificially illuminated room, completely free from chemical contamination. They were fed with the standard laboratory food and allowed to access it and drink water freely. All rats received humane care in compliance with the Ethical Committee of Zagazig University and in accordance with the NIH Guidelines for the Care and Use of Laboratory Animals. **I confirm that the experimental protocol was approved by Zoology department-Faculty of science-Zagazig University**.

### 2.2 Experimental design

The animals were assigned to 3 groups:

**Group I (control group)**: included ten rats that received no treatment for 2 months.

**Group II (DM group)**: included 10 rats received dimethoate (20 mg/kg body weight) dissolved in 1ml corn oil [1/20 of the LD50 (380 mg/kg)] [14] once daily for 14 weeks via oral intubation.

**Group III (DM+ALA group)**: included 10 rats received dimethoate (20 mg/kg body weight) dissolved in 1ml corn oil [1/20 of the LD50 (380 mg/kg)] once daily for 14 weeks via oral intubation [14] then received ALA (50 mg/kg) [15]

At the end of the experiment, Blood samples were collected [16], and then rats of all groups were sacrificed with the intraperitoneal injection of 25 mg/ kg sodium thiopental [17]. Liver and kidney sections were taken and fixed. Samples were processed for light microscope examination.

### 2.3 Experimental procedures

Dimethoate (purity 98%), Nacetylcysteine were purchased from Sigma Chemicals, St. Louis, MO, USA. Alphalipoic acid capsule (thiotex forte 600 mg/capsule): it was obtained from Marcryrl pharmaceutical industries, El-obour city –Egypt.

#### 2.3.1 Collection of blood samples

Blood samples were collected from the retro-orbital venous plexus, under mild anesthesia, using a fine heparinized capillary tube introduced into the medial epicanthus of the rat’s eye [72]. Blood samples were collected in a clean graduated centrifuge tube, allowed to clot at room temperature for 10 min and then centrifuged using Remi cool centrifuge at 3000 rpm for 20 min. The supernatant serum was collected in a dry clean tube for estimation of biochemical parameters. All biochemical procedures were conducted in the Central Research Lab, Faculty of medicine, Zagazig University.

#### 2.3.2 Biochemical estimation of serum urea, creatinine and BUN

Serum urea was estimated by quantitative colorimetric method using QuantiChrom^TM^ assay kits (BioAssay Systems, USA). The BUN was measured using a commercial kit (BUN II reagent kit; Wako Pure Chemical Industries) [18].

#### 2.3.3 Biochemical estimation of serum lipid profile

Triglycerides, total cholesterol, HDL-cholesterol and LDL-cholesterol concentration were evaluated enzymatically using assay kits (Sigma Chemical Co, St Louis, MO, USA). VLDL-cholesterol was calculated as triglycerides and LDL-cholesterol was calculated by the equation: LDL-cholesterol=Total serum cholesterol – (HDL+VlDl) [19].

#### 2.3.4 Biochemical estimation of serum ALT, AST, ALP and TP

The activities of serum ALT, ALP and AST were determined spectrophotometrically using an Automated analyzer method (Opera Technician Bayer Auto analyzer). Serum total protein (TP) Level was determined by colorimetric point method.

#### 2.3.5 Histopathological study

All steps were conducted at Histology and Cell Biology Department, Faculty of Medicine, Zagazig University. Specimens from the liver and renal cortex of each animal were fixed in 10% neutral formol saline, dehydrated, embedded in paraffin wax and processed into 5 μm thick sections. They were stained with hematoxylin and eosin. Stained slides were analyzed by light microscopy in Image Analysis Unit [20].

#### 2.3.6 Statistical analysis

Data were recorded and entered using the statistical package SPSS version 13. Data were described using mean and standard error for quantitative variables. Comparisons between groups were done using one-way analysis of variance with multiple comparisons post hoc test [21]. Results were considered statistically significant at P < 0.05, while P values less than 0.001 were highly significant.

## 3. Results

### 3.1 Biochemical Results

#### Biochemical Estimation serum Liver functions

DM-treated group revealed a high significantly increasing in mean of ALT, AST and ALP compared to control group (P < 0.01) while it showed high significant decreasing in mean of TP. DM+ALA group publicized a high significantly decreasing in mean of ALT, AST and ALP (P < 0.01) compared to DM-treated group, while it showed a significant increasing in mean of TP (P<0.05). Statistical results of various studied groups are presented in (**Table 1**).

**Table 1:** Serum TP, ALT, AST and ALP in various studied groups.

|  |  | Control | Dimethoate (DM) | DM+ALA |
| --- | --- | --- | --- | --- |
| T. protein (TP)<br>(g/dl) | Mean±SE | 6.76±0.19 | 5.32±0.37 | 6.51±0.28 |
|  | % change |  | -21.30<sup>a</sup> | +22.36<sup>c</sup> |
|  | Significance |  | <0.01**<sup>a</sup> | <0.05*<sup>c</sup> |
| ALT<br>(IU/L) | Mean±SE | 15.39±1.02 | 23.72±1.52 | 16.53±0.74 |
|  | % change |  | +54.13<sup>a</sup> | -30.31<sup>c</sup> |
|  | Significance |  | <0.01**<sup>a</sup> | <0.001**<sup>c</sup> |
| AST<br>(IU/L) | Mean±SE | 16.56±0.94 | 21.77±1.39 | 15.65±.26 |
|  | % change |  | +31.46<sup>a</sup> | -28.11<sup>c</sup> |
|  | Significance |  | <0.01**<sup>a</sup> | <0.01**<sup>c</sup> |
| ALP<br>(IU/L) | Mean±SE | 70.48±1.84 | 120.97±6.64 | 76.73±4.56 |
|  | % change |  | +71.64<sup>a</sup> | -36.57<sup>c</sup> |
|  | Significance |  | <0.01**<sup>a</sup> | <0.01**<sup>c</sup> |

\*P<0.05: Significance, \*\*P<0.01: High significance & P > .05:Non-Significance
<sup>a</sup> is comparison of control vs DM, <sup>c</sup> is comparison of DM vs DM+ALA

#### Biochemical estimation of serum urea, creatinine, BUN and testosterone

**Table (2)** showed that DM-treated group demonstrated a high significant increase in mean of Urea compared to control group (P<0.01) while it showed a significant increase (P<0.05) in mean of creatinine and a a high significant decrease in testosterone level. DM+ALA group shown a high significantly decreasing in mean of Urea, creatinine and BUN (P<0.01) compared to DM-treated group, while it showed a significant increasing in mean of testosterone level (P<0.01).

**Table 2:** Serum Urea, Creatinine, BUN and Testosterone in various studied groups.

|  |  | Control | Dimethoate (DM) | DM+ALA |
| --- | --- | --- | --- | --- |
| Urea (mg/dl) | Mean±SE | 56.67±2.55 | 83.89±1.85 | 48.22±3.43 |
|  | % change |  | +48.03<sup>a</sup> | -42.51<sup>c</sup> |
|  | Significance |  | <0.01**<sup>a</sup> | <0.01**<sup>c</sup> |
| Creatinine (mg/dl) | Mean±SE | 0.63±0.009 | 0.79±0.04 | 0.63±0.60 |
|  | % change |  | +25.39<sup>a</sup> | -20.25<sup>c</sup> |
|  | Significance |  | <0.05*<sup>a</sup> | <0.01**<sup>c</sup> |
| BUN (mg/dl) | Mean±SE | 25.94±1.14 | 26.88±1.48 | 19.97±1.05 |
|  | % change |  | +3.62<sup>a</sup> | -25.70<sup>c</sup> |
|  | Significance |  | > .05<sup>a</sup> | <0.01**<sup>c</sup> |
| Testosterone (nM) | Mean±SE | 4.42±0.12 | 3.48±0.09 | 4.35±0.08 |
|  | % change |  | -21.26<sup>a</sup> | 25<sup>c</sup> |
|  | Significance |  | <0.01**<sup>a</sup> | <0.01**<sup>c</sup> |

\*P<0.05: Significance, \*\*P<0.01: High significance & P > 0.05: Non-Significance
<sup>a</sup> is comparison of control vs DM, <sup>c</sup> is comparison of DM vs DM+ALA

#### Biochemical Estimation serum Lipid profile

DM-treated group exhibited statistically significantly increasing in mean of TG, LDL-c, VLDL-c and Atherogenic index compared to control group (P < 0.05) while it showed non-significance decreasing mean in TC and HDL-c. DM+ALA group revealed non-significant lower mean TG, LDL-c, and Atherogenic index compared to DM-treated group (P > 0.05). Moreover, statistically significant increase of TC (P < 0.05) but non-significant increase of HDL-c and VLDL-c (P > 0.05). Statistical results of various studied groups are presented in **(Table 3).**

**Table 3:** Serum Lipid profile in various studied groups.

|  |  | Control | Dimethoate (DM) | DM+ALA |
| --- | --- | --- | --- | --- |
| TC (mg/dl) | Mean±SE | 151.01±6.30 | 150.25±9.14 | 178.37±4.78 |
|  | % change |  | -0.50<sup>a</sup> | +18.71<sup>c</sup> |
|  | Significance |  | > .05<sup>a</sup> | <0.05*<sup>c</sup> |
| TG (mg/dl) | Mean±SE | 183.25±2.92 | 211.37±12.06 | 192.62±10.85 |
|  | % change |  | +15.34<sup>a</sup> | -8.87<sup>c</sup> |
|  | Significance |  | <0.05*<sup>a</sup> | > .05<sup>c</sup> |
| HDL-C (mg/dl) | Mean±SE | 54.75±2.36 | 44.50±12.05 | 50.62±2.37 |
|  | % change |  | -18.72<sup>a</sup> | +13.75<sup>c</sup> |
|  | Significance |  | > .05<sup>a</sup> | > .05<sup>c</sup> |
| LDL-C (g/dl) | Mean±SE | 161.76±10.15 | 200.23±9.04 | 182.25±2.05 |
|  | % change |  | +23.78<sup>a</sup> | -8.97<sup>c</sup> |
|  | Significance |  | <0.05*<sup>a</sup> | > .05<sup>c</sup> |
| VLDL-C (mg/dl) | Mean±SE | 34.77±0.70 | 42.35±2.66 | 43.51±2.30 |
|  | % change |  | +21.80<sup>a</sup> | +2.73<sup>c</sup> |
|  | Significance |  | <0.05*<sup>a</sup> | > .05<sup>c</sup> |
| Atherogenic index | Mean±SE | 0.183±0.02 | 0.253±0.02 | 0.224±0.02 |
|  | % change |  | +38.25<sup>a</sup> | -11.46<sup>c</sup> |
|  | Significance |  | <0.05*<sup>a</sup> | > .05<sup>c</sup> |

\*P<0.05: Significance, \*\*P<0.01: High significance & P > 0.05:Non-Significance
<sup>a</sup> is comparison of control vs DM , <sup>c</sup> is comparison of DM vs DM+ALA

### 3.2 Histopathological Results

**Fig. 1:**
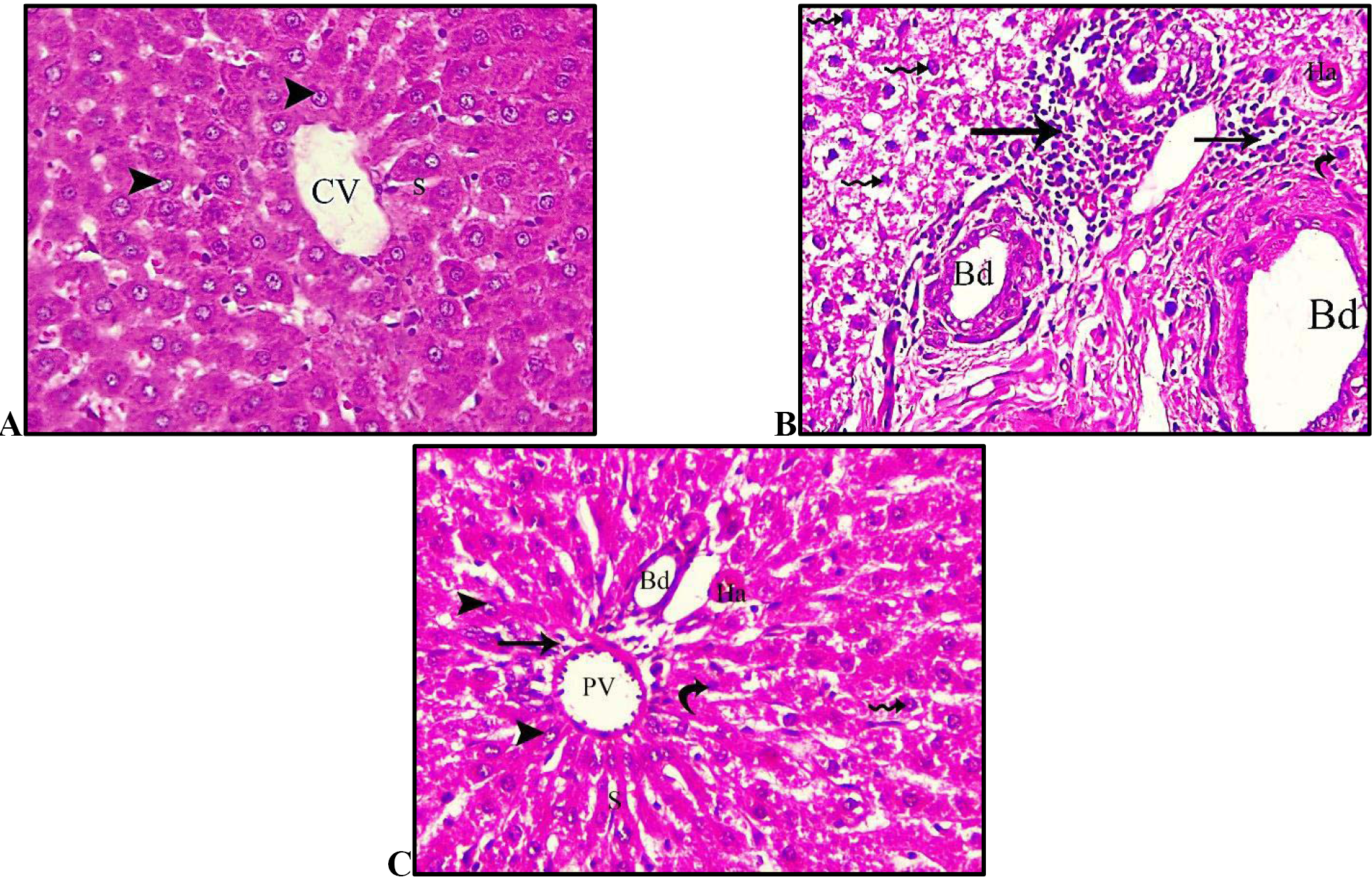
photomicrographs of sections in the liver of adult male rats of different groups. **(A): control group** showing normal hepatic laminae, hepatocytes and intact nuclei **(arrowheads).** Central vein **(CV)** lies at the centre of the lobule surrounded by the hepatocytes strands (with strongly eosinophilic granulated cytoplasm) and separated by blood sinusoids **(S). (B): DM group** showing focal necrosis with fibrosis and inflammatory infiltration **(arrow).** Also it is noticed a congestion of hepatic sinusoids and dilated bile duct **(Bd),** Hepatocytes lose its polygonal architecture and have a vacuolated cytoplasm with deeply stained pyknotic nuclei (zigzag arrow) while heaptic artery **(Ha)** is congested and apoptotic bodies **(curved arrow)** also appeared. **(C): DM+ALA group** showing a restoring of normal architicture of the hepatic strands, hepatocytes and normal nuclei **(arrowheads).** Portal vein **(PV)** and Bile duct **(Bd)** appeared with normal shape while apoptotic bodies **(curved arrow)** highly decreased.

**Fig. 2:**
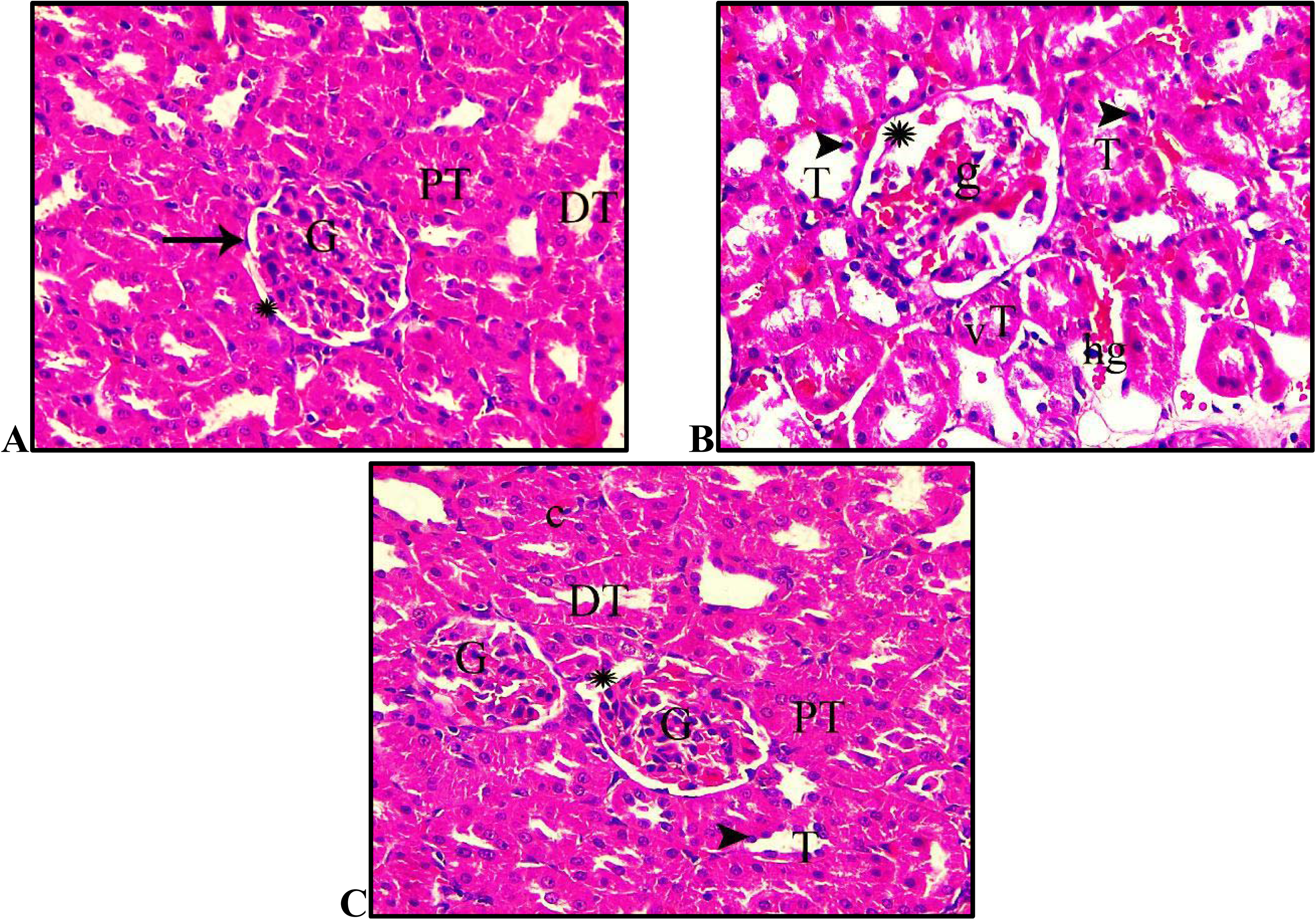
photomicrographs of sections in the renal cortex of adult male rats of different groups. **(A): control group** showing normal glomeruli **(G)**, Bowman’s capsules lined by intact simple squamous cells **(arrow)**, narrow Bowman’s space **(star)**, proximal convoluted tubules PCT with cuboidal epithelium and narrow lumen **(PT)**, and distal convoluted tubules DCT wide lumen **(DT)**. **(B): DM group showing** shrunken and segmented glomerulus **(g)**, marked hemorrhage **(hg)** at distal convoluted tubules and wide Bowman’s spaces **(star)**. Renal tubules **(T)** reveal exfoliated cells with dark-stained apoptotic nuclei **(arrowhead)** and vacuolation **(V)** of renal tubule cells. **(C): DM+ALA group** showing nearly normal glomeruli **(G)** but with wide Bowman’s spaces **(star).** Some tubule**s (T)** appear with darkly stained nuclei **(arrowhead)** and capillaries between tubules **(c)**. Normal proximal convoluted tubules **(PT)**, distal convoluted tubules **(DT)** are seen.

## 4 Discussion

DM mediated oxidative stress is responsible for the inhibition of cholinesterase activity [22]. According to [23], dimethoate sub chronic exposure has led to a change in the total antioxidant status in the liver, brain and kidney of rats. After dimethoate exposure there was an increase in lipid peroxidation, catalase, and glutathione reductase and superoxide dismutase activities [24]. DM affects the functions of multiple organs including liver. Dimethoate was reported to alter the level of the marker parameters related to the liver in rats and mice [6]. Numerous studies indicate that dimethoate intoxication can cause oxidative stress by the generation of free radicals and induce hepatic lipid peroxidation in mice [25] and rats [14].

Therefore, it is reasonable to assume that the significant elevation in the levels of the liver enzymes (ALT, AST and ALP) was due to the administration of DM for (20 mg/kg b.wt.) for 14 weeks as shown in **table (1)**. These results came in the same line with [26] who recorded a marked increase in the level of alanin-aminotransferse (ALT) asparatatea-aminotransferase (AST) and the level of alkaline phosphatase (ALP) in the serum of Dimethoate treated group. Also, Alteration in protein metabolic profiles was dose – and time- dependent. According to [24] there was decrease in serum total protien, albumin and globulin and increase in bilirubin level in dimethoate treated rats.

The reduction in serum protein could be attributed to changes in protein and free amino acid metabolism and their synthesis in the liver [27]. Also, the protein level suppression may be due to loss of protein either by reducing in protein synthesis or increased proteolytic activity or degradation [28]. In the light of current knowledge, the present results showed a significant lowering in the total protein levels after treatment of the rats with DM. This finding was confirmed by [24] who found that DM decrease serum total protein and albumin. Oxidative stress can be viewed as the disorder in the oxidant-antioxidant balance in favor of the past.

Histopathological results revealed a great impairment of DM on the architecture of the hepatocytes and liver constituents, where degenerated hepatocytes, pyknotic nuclei, focal fibrosis, dilated biliary ducts and inflammatory infiltrations are appeared in DM group as illustrated in **fig.(4)**. These observations are in agreement with [29] who elucidated that mice exposed to dimethoate showed a degenerative changes in the liver including congestion blood vessels, infiltration, vasodilatation and hydropic changes. Also these investigations goes with [41] who clarified that DM caused the following in the liver lymphocytic infiltration, congestion, nuclear death, enlargement of hepatic sinusoids, hepatocellular damage, cytoplasmic vacuolation and degeneration in nuclei. This is due to changes in cellular integrity and membrane permeability due to exposure to toxic chemicals [30]. Also DM being lipophilic substance can interact with cellular membranes [31] and the membrane damage or necrosis releases such enzymes into the circulation [30]. While ALA giving caused a restoring of normal architecture of the hepatic strands, hepatocytes and normal nuclei. Portal vein and Bile duct appeared with normal shape while apoptotic bodies highly decreased.

Numerous animal studies have reported that ALA has blood lipid modulating properties beyond its antioxidant and anti-inflammatory features [32]. ALA has been affirmed to have various gainful impacts, avoiding and treating many diseases through its antioxidant and anti-inflammatory activities [33]. ALA is a disulfide compound that functions as a coenzyme in pyruvate dehydrogenase and alpha–ketoglutarate dehydrogenase mitochondrial reactions, leading to the production of cellular energy (ATP) [34]. The protective role of ALA against hepatotoxicity and oxidative stress is well documented in a series of scientific reports [35]. Our results showed that administration of ALA to treated rats with DM caused a significant recovery to the biochemical marker of the liver where ALT, AST and ALP are approximately returned to the normal levels as explained in **table (1)**. These results were in line with [36] who proved that adding ALA to the diet of rats receiving a high-fat diet resulted in the noticeable reduction in ALT and AST. These findings are interpreted by [37] who reported that ALA and its reduced form, dihydrolipoic acid, reduce oxidative stress by scavenging a number of free radicals in both membrane and aqueous areas by stopping membrane lipid peroxidation and protein impairment through the redox regeneration of other antioxidants such as vitamins C and E, and by elevating intracellular glutathione. In addition to its role as an antioxidant, ALA can affect the activity of enzymes at various levels of metabolic pathways. Also, [38] showed that ALA can affect mammalian pyruvate dehydrogenase complex (PDC) based on its stereo-selectivity. It has been shown that short-term administration of ALA at high dosage to normal rats caused an inhibition of gluconeogenesis secondary to an interference with hepatic fatty acid oxidation [39], and increased plasma levels of pyruvate and lactate were observed in ALA treatments [40].

[41] stated that creatinine is an amino acid produced as a waste product of creatine, which acts as an important energy storage in muscle metabolism. Urea and creatinine level in blood rises when there is kidney impairment which prevents the kidneys from filtering urea and creatinine out of the blood. [41] also suggested significant increase in urea and creatinine level in serum after dimethoate exposures for 15 and 30 days. These investigations suggested that DM induced hepatic injury and also indicate that kidneys may suffer to clear the waste products and toxins from the blood [4]. Interestingly, our results approved that the levels of urea, Creatinine and BUN are increased significantly in the DM group due to induction of oxidative effects by DM as shown in **table (2)**. Organophosphorus induces H2O2 production and lipid peroxidation in a kidney cell [42]. Others have provided additional evidence for the occurrence of Organophosphorus-induced oxidative tissue damage evidenced by DNA-strand breaks, increased activities of anti- oxidant enzymes [43] and down-regulation of glutathione peroxidase activity and glutathione [44]. Lipid peroxidation, ATP depletion, DNA damage, protein oxidation, and intra- cellular calcium increase due to membrane permeability lesions may be the pathway for renal cell damage by organophosphorus [45]. In the present study, ALA administration caused a great lowering in the mean of Urea, creatinine and BUN, while it showed a significant increasing in mean of testosterone level as shown in **table (2)**. It is agreed with An animal model has shown that ALA protects against ischemic acute renal failure, as exemplified by attenuation of blood urea nitrogen (BUN), plasma concentration of creatinine, urinary osmolality, creatinine clearance (SCr), and fractional excretion of Na+, as well as attenuation of tubular necrosis, proteinaceous casts and medullary congestion in the renal tissue,) the protective effects may relate in part to decreased content of endothelin-1 (ET-1) in the kidney [46]. ALA resulted in increases in total antioxidant capacity and reduced GSH and Na+/K+-ATPase activity [47].

Histopathological results of the kidney explained that DM administration to rats caused destructive alterations in the architecture of kidney where DM caused shrunken and segmented glomerulus, marked hemorrhage at distal convoluted tubules and wide Bowman’s spaces. Also, renal tubules reveal exfoliated cells with dark-stained apoptotic nuclei and vacuolation of renal tubule cells as observed in **fig. (2)**. These alterations are confirmed by [29] who observed that DM caused Histological changes in the Kidney including Glomerular Degeneration, Tubular Degeneration, Hemorrhage, Infiltration, Hydropic Changes, Tubular Cast, Tubular Widened Lumen and Glomerular Shrinkage. While in ALA group nearly normal glomeruli restored but with wide Bowman’s spaces. Also, Normal proximal convoluted tubules, distal convoluted tubules are seen. It is confirmed by ALA regenerated and reduced tubular dilation and regenerated tubular epithelium [47]. It was also found that α-LA prevented eNOS and neuronal NOS, but decreased inducible NOS. Meanwhile, ALA decreased expression levels of ET-1 [35]. Both in vitro and in vivo studies revealed that ALA reduced serum levels of BUN and SCr, decreased MDA levels and ROS levels as well as alleviated the severity of kidney lesions (tubular cell necrosis, cytoplasmic vacuolation, hemorrhage and tubular dilatation) [48].

The current research illustrated that DM group exhibited a significant increasing in TG, LDL-c, VLDL-c and Atherogenic index compared to control group while it showed non-significance decreasing mean in TC and HDL-c as in **table (3)**. It is partially accepted by [49] who stated that DM caused an increase in the serum total cholesterol level. Increased serum cholesterol can be attributed to the effects of the pesticide on the permeability of liver cell membranes [50]. Also, the increase in the level of serum total cholesterol may be attributed to the blockage of the liver bile ducts, causing a reduction or cessation of cholesterol secretion into the duodenum [51]. An increase in the serum cholesterol level may be a sign of liver damage [52]. Also, [49] demonstrated that DM caused decreases in the triglyceride and VLDL-cholesterol levels. Some different pesticides cause a decrease in the VLDL-cholesterol and triglyceride levels [53].

Apart from oxidative stress generated by DM, there was also a significant reduction in the level of plasma HDL along with increment in the levels of triglyceride, LDL and total cholesterol was noted in DM group. Free radicals enhance the oxidation of LDL and oxidized LDL affects many biological processes involved in atherogenesis. High HDL levels competes with LDL receptors site on the smooth muscle cells of arterial wall and inhibits the uptake of LDL and thus prevents the organism from the risk of atherogenesis [54]. In ALA group TG, LDL-c, and Atherogenic index are declined and approximately returned to the normal levels of the control group but an increasing of HDL-c and VLDL-c took place. It is in line with [55] who clarified that ALA caused a significant increase in HDL with a concurrent significant reduction in TC, TG, LDL, and VLDL (bad cholesterol). Moreover, [56] reported decreased serum TC, TG, and LDL-C levels in obese subjects, following ALA supplementation. The probable mechanism for theses alterations is increasing of insulin sensitivity and controlled activities of the enzymes involved in lipolysis and triglyceride synthesis [57]. Furthermore, ALA appears to increase activities of lipoprotein lipase as well as Lecithin Cholesterol Acyltransferase (LCAT) [58].

## Conclusion

It is concluded that ALA therapy can ameliorate the negative effects of DM that affect the vital organs as the liver and kidney. Also ALA can reduce the occurrence of atherogensis by reducing the levels of bad cholesterol in the blood. ALA boosts the levels of testosterone so it augments the male sexual characters.

## Abbreviations

DM: Dimethoate
ALA: Alpha-lipoic acid
BUN: Blood urea nitrogen
TC: total cholesterol
LDL: Low density lipoprotein
HDL: High density lipoprotein
TG: Triglycerides
Ops: organophosphates
and DHLA: Dihydrolipoic acid

## Declarations

### Acknowledgment

To who installed in me the value of education and the rewards and opportunities it can generate to our parents especially my Father who supplied me with enthusiasm, support and creative insight. His critical reading of the manuscript that helped me in refining the concept of this thesis, his deep interest in the topic and unfailing encouragement are highly appreciated.

### Ethical approval and consent to participate

All applicable international, national, and/or institutional guidelines for the care and use of animals were followed.

### Availability of data materials

’Not applicable’

### Author contributions

HA carried out the physiological, histological, biochemical and anatomical studies, participated in the sequence alignment and drafted the manuscript.

---

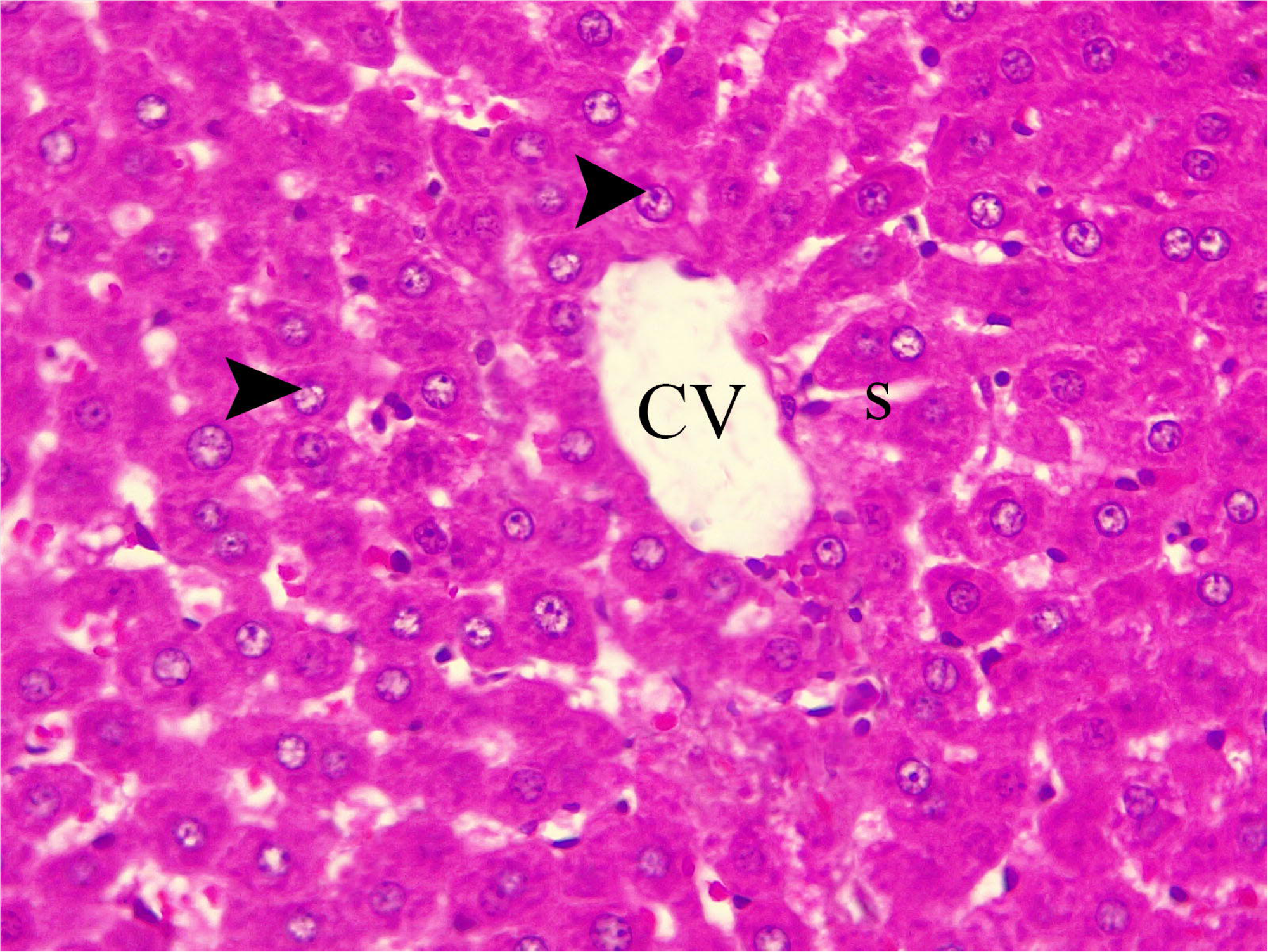

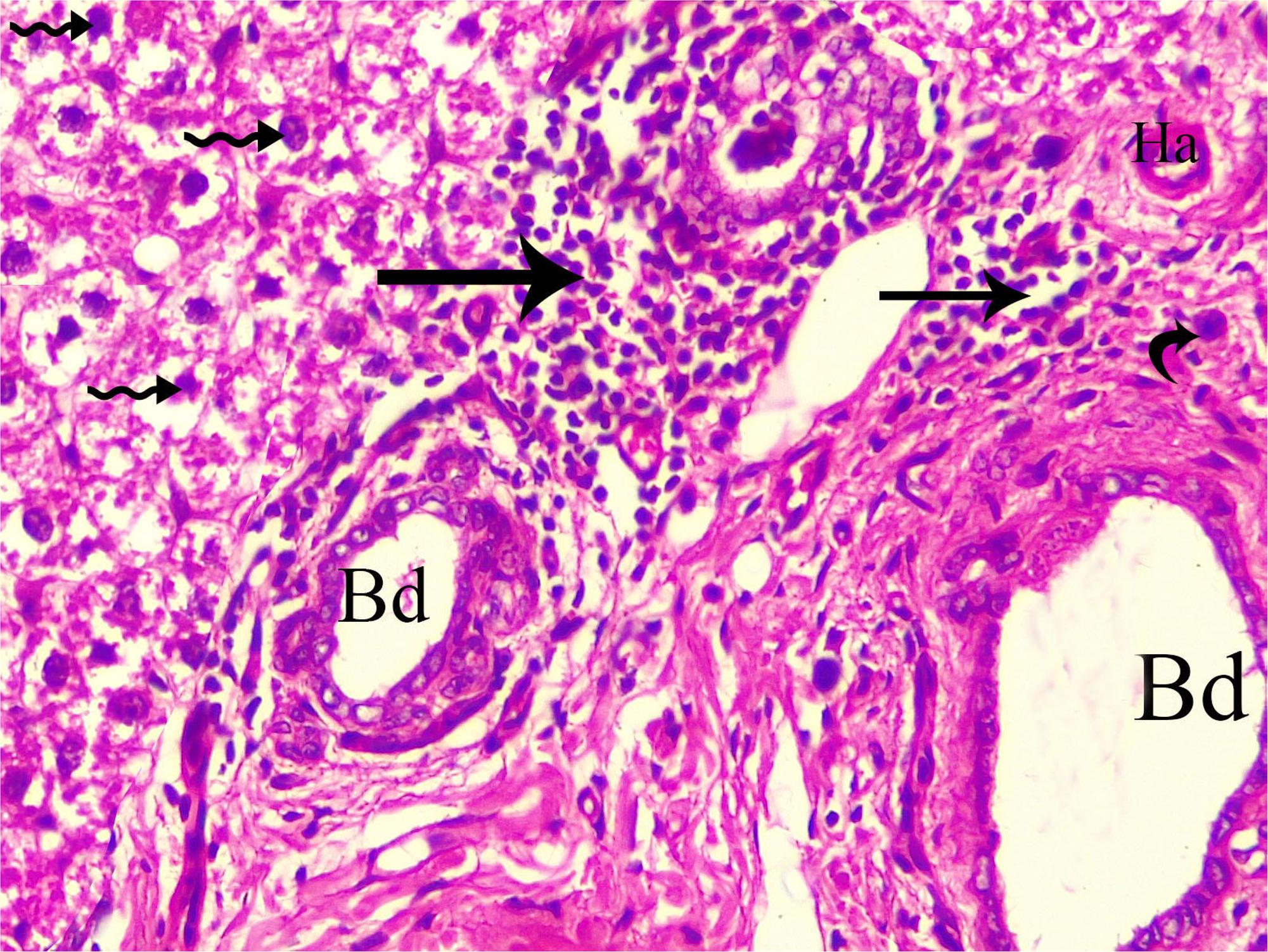

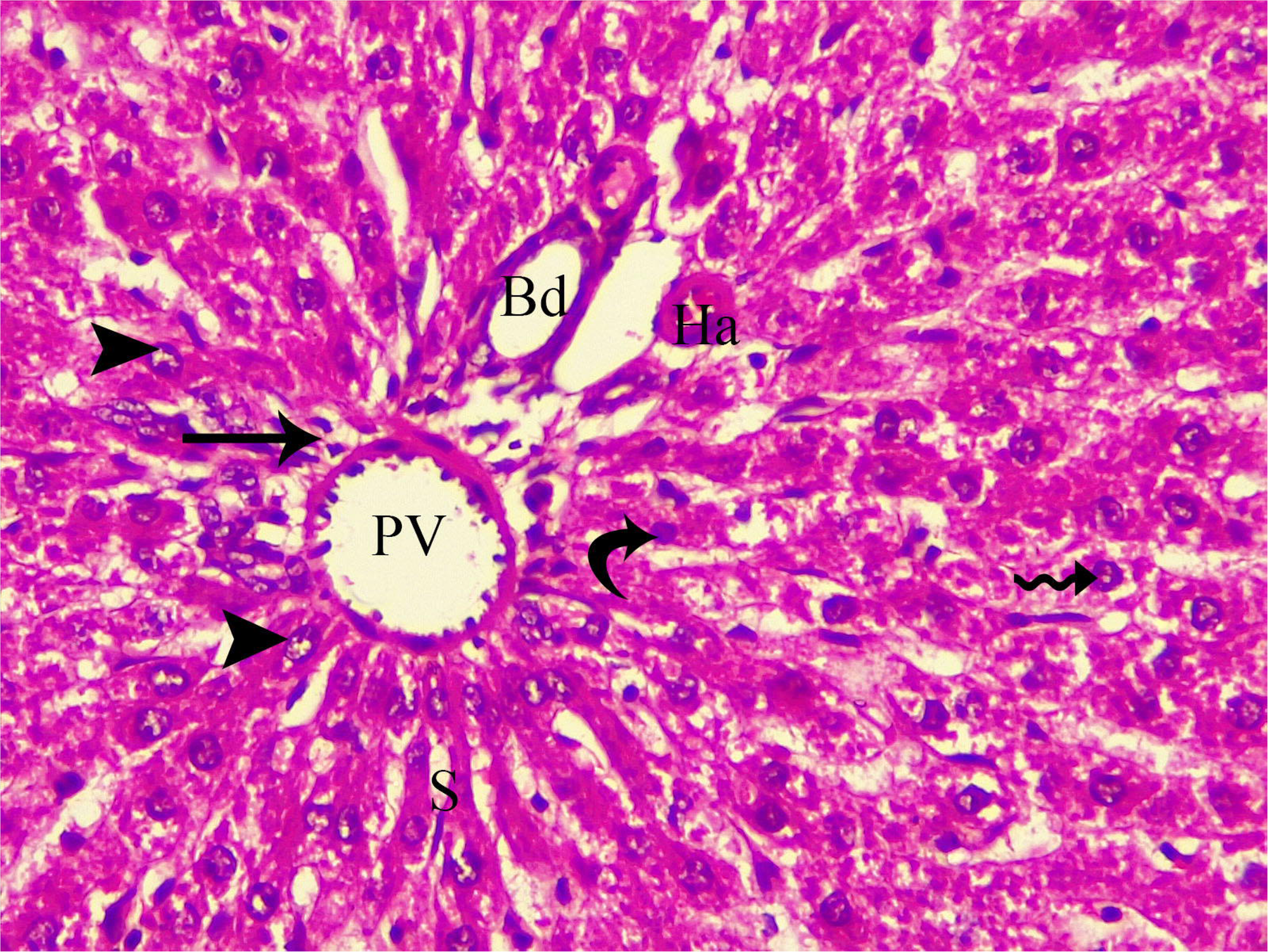

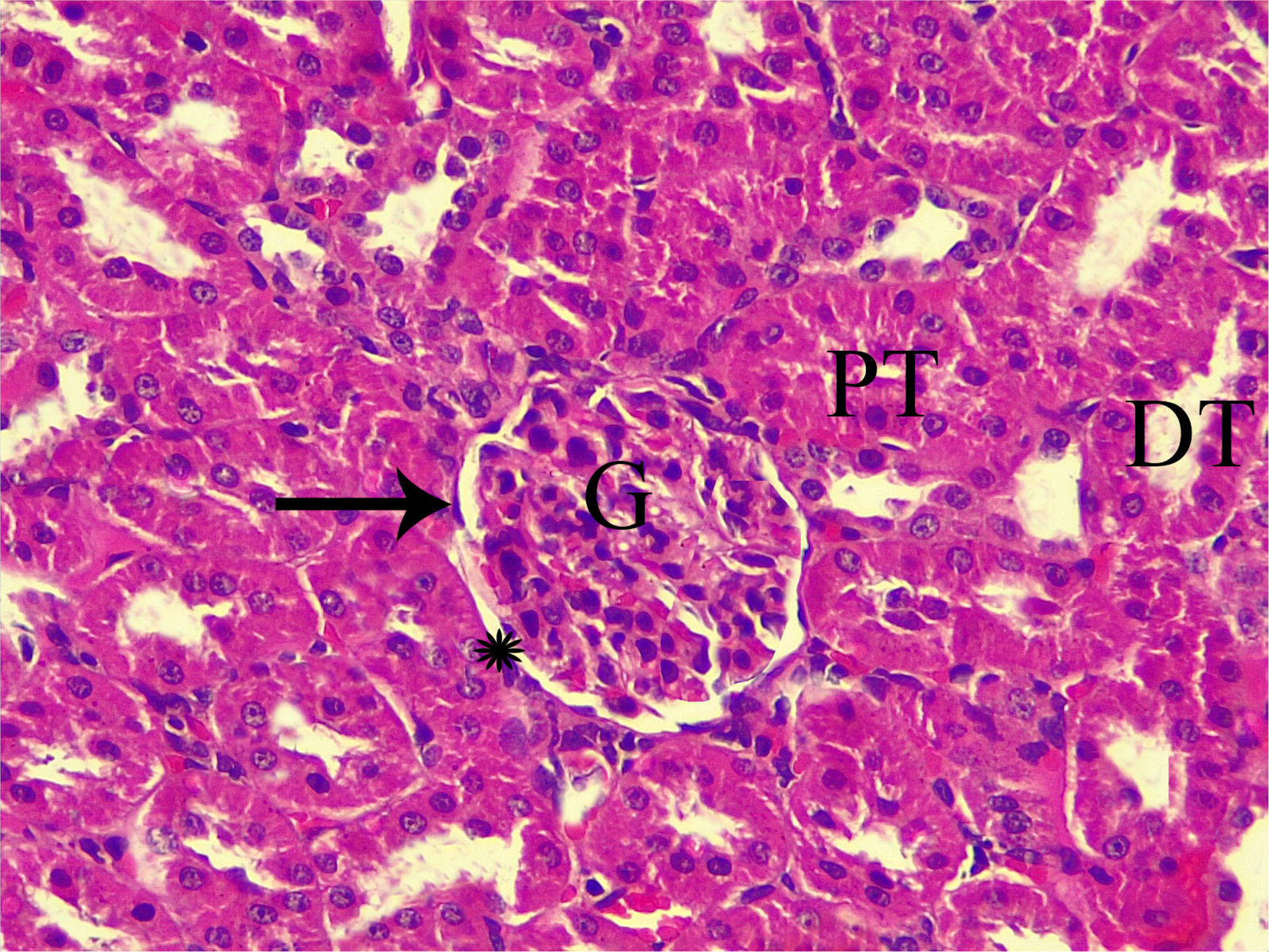

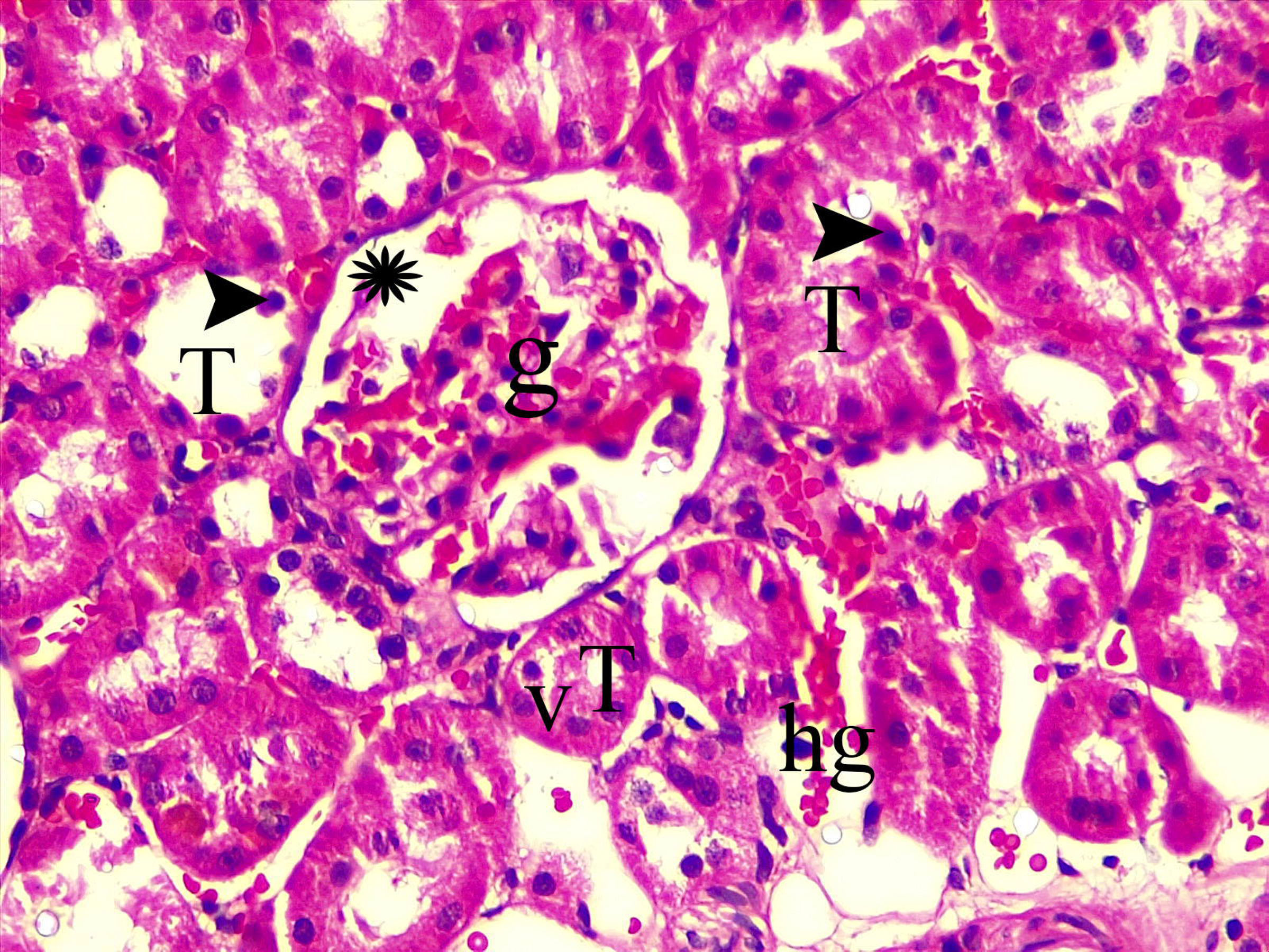

## References

1. Lukaszewicz-Hussain, A., Role of oxidative stress in organophosphate insecticide toxicity–Short review. Pesticide biochemistry and physiology, 2010. 98(2): p. 145–150.

2. Nazam, N., et al., Combined in silico and in vivo studies shed insights into the acute acetylcholinesterase response in rat and human brain. Biotechnology and applied biochemistry, 2015. 62(3): p. 407–415.

3. Shadnia, S., et al., Evaluation of oxidative stress and genotoxicity in organophosphorus insecticide formulators. Human & experimental toxicology, 2005. 24(9): p. 439–445.

4. Amara, I.B., et al., Dimethoate induced oxidative damage and histopathological changes in lung of adult rats: modulatory effects of selenium and/or vitamin E. Biomedical and Environmental Sciences, 2012. 25(3): p. 340–351.

5. Ogutcu, A., et al., The effects of organophosphate insecticide diazinon on malondialdehyde levels and myocardial cells in rat heart tissue and protective role of vitamin E. Pesticide biochemistry and physiology, 2006. 86(2): p. 93–98.

6. Khan, A.A., M.A. Shah, and S.U. Rahman, Occupational exposure to pesticides and its effects on health status of workers in Swat, Khyber Pakhtunkhwa, Pakistan. Journal of Biology and Life Science, 2013. 4(2): p. 43–55.

7. Crissman, J.W., et al., Best practices guideline: toxicologic histopathology. Toxicologic Pathology, 2004. 32(1): p. 126–131.

8. Gorąca, A., et al., Lipoic acid–biological activity and therapeutic potential. Pharmacological Reports, 2011. 63(4): p. 849–858.

9. Singh, U. and I. Jialal, Retracted: Alpha-lipoic acid supplementation and diabetes. Nutrition reviews, 2008. 66(11): p. 646–657.

10. Wongmekiat, O., D. Leelarungrayub, and K. Thamprasert, Alpha-lipoic acid attenuates renal injury in rats with obstructive nephropathy. BioMed research international, 2013. 2013.

11. Kang, K.P., et al., Alpha-lipoic acid attenuates cisplatin-induced acute kidney injury in mice by suppressing renal inflammation. Nephrology Dialysis Transplantation, 2009. 24(10): p. 3012–3020.

12. Bilska, A. and L. Wlodek, Lipoic acid-the drug of the future. Pharmacol Rep, 2005. 57(5): p. 570–577.

13. Ayhan, M., et al., PREVENTİVE EFFECTS OF ALPHA-LİPOİC ACİD ON DİABETİC NEPHROPATHY İN A RAT MODEL. ACTA MEDICA MEDITERRANEA, 2014. 30(6): p. 1221–1225.

14. Kamath, V., A.K.R. Joshi, and P. Rajini, Dimethoate induced biochemical perturbations in rat pancreas and its attenuation by cashew nut skin extract. Pesticide biochemistry and physiology, 2008. 90(1): p. 58– 65.

15. Emam, A.M., et al., Protective effects of alpha-lipoic acid and coenzyme Q10 on lipopolysaccharide-induced liver injury in rats.

16. Wang, S.-J., et al., Mechanism of treatment of kidney deficiency and osteoporosis is similar by Traditional Chinese Medicine. Current pharmaceutical design, 2016. 22(3): p. 312–320.

17. Kara, A., et al., Ultra-structural changes and apoptotic activity in cerebellum of post-menopausal-diabetic rats: a histochemical and ultra-structural study. Gynecological Endocrinology, 2014. 30(3): p. 226–231.

18. Roubalová, L., et al., Flavonolignan 2, 3-dehydrosilydianin activates Nrf2 and upregulates NAD (P) H: quinone oxidoreductase 1 in Hepa1c1c7 cells. Fitoterapia, 2017. 119: p. 115–120.

19. Chattopadhyay, R. and M. Bandyopadhyay, Effect of Azadirachta indica leaf extract on serum lipid profile changes in normal and streptozotocin induced diabetic rats. African Journal of Biomedical Research, 2005. 8(2): p. 101–104.

20. Bancroft, J.D. and C. Layton, The hematoxylins and eosin. Bancroft’s Theory and Practice of Histological Techniques. Elsevier, 2013: p. 173–186.

21. Armitage, P., G. Berry, and J.N.S. Matthews, Statistical methods in medical research. 2008: John Wiley & Sons.

22. Bai, Y., L. Zhou, and J. Wang, Organophosphorus pesticide residues in market foods in Shaanxi area, China. Food Chemistry, 2006. 98(2): p. 240–242.

23. Tarbah, F., et al., Distribution of dimethoate in the body after a fatal organphosphateintoxication. Forensic science international, 2007. 170(2-3): p. 129–132.

24. Attia, A.M. and H. Nasr, Dimethoate-induced changes in biochemical parameters of experimental rat serum and its neutralization by black seed (Nigella sativa L.) oil. Slovak Journal of Animal Science (Slovak Republic), 2009.

25. Sivapiriya, V. and S. Venkatraman, Effects of dimethoate (O, O-dimethyl S-methyl carbamoyl methyl phosphorodithioate) and ethanol in antioxidant status of liver and kidney of experimental mice. Pesticide biochemistry and physiology, 2006. 85(2): p. 115–121.

26. El-Damaty, E., et al., Biochemical and histopathological effects of systemic pesticides on some functional organs of male albino rats. J. Appl. Sci. Res, 2012. 8(11): p. 5459–5469.

27. Salim, A., et al., Influence of pomegranate (Punica granatum L.) on dimethoate induced hepatotoxicity in rats. Int J Biol Biomol Agr Food Biotechn Eng, 2014. 8(8): p. 908–913.

28. Yeragi, S., A. Rana, and V. Koli, Effect of pesticides on protein metabolism of mud skipper Boleophthalmus dussumieri. Journal of Ecotoxicology & Environmental Monitoring, 2003. 13(3): p. 211– 214.

29. Alarami, A.M., Histopathological Changes in the Liver and Kidney of Albino Mice on Exposure to Insecticide, Dimethoate. Int. J. Curr. Microbiol. App. Sci, 2015. 4(7): p. 287–300.

30. Ajani, E., et al., Toxicological implications of continuous administration of aqueous leaves extract of hydrocotyl bonariensis in rats. Archives of Applied Science Research, 2011. 3(5): p. 471–478.

31. Saafi, E.B., et al., Protective effect of date palm fruit extract (Phoenix dactylifera L.) on dimethoate induced-oxidative stress in rat liver. Experimental and Toxicologic Pathology, 2011. 63(5): p. 433–441.

32. Zulkhairi, A., et al., Alpha lipoic acid posses dual antioxidant and lipid lowering properties in atherosclerotic-induced New Zealand White rabbit. Biomedicine & Pharmacotherapy, 2008. 62(10): p. 716–722.

33. Odabasoglu, F., et al., α-Lipoic acid has anti-inflammatory and anti-oxidative properties: an experimental study in rats with carrageenan-induced acute and cotton pellet-induced chronic inflammations. British Journal of Nutrition, 2011. 105(1): p. 31–43.

34. Hamzawy, M.A., et al., Hepatoprotective effect of estradiol and-Lipoic Acid in Rats. Global Journal of Pharmacology, 2014. 8(4): p. 694–702.

35. Bae, E.H., et al., Effects of α-lipoic acid on ischemia-reperfusion-induced renal dysfunction in rats. American journal of physiology-renal physiology, 2008. 294(1): p. F272–F280.

36. Valdecantos, M.P., et al., Lipoic acid administration prevents nonalcoholic steatosis linked to long-term high-fat feeding by modulating mitochondrial function. The Journal of nutritional biochemistry, 2012. 23(12): p. 1676–1684.

37. Evans, J.L. and I.D. Goldfine, α-Lipoic acid: a multifunctional antioxidant that improves insulin sensitivity in patients with type 2 diabetes. Diabetes technology & therapeutics, 2000. 2(3): p. 401–413.

38. Hong, Y.S., et al., The inhibitory effects of lipoic compounds on mammalian pyruvate dehydrogenase complex and its catalytic components. Free Radical Biology and Medicine, 1999. 26(5-6): p. 685–694.

39. Khamaisi, M., et al., Lipoic acid acutely induces hypoglycemia in fasting nondiabetic and diabetic rats. Metabolism, 1999. 48(4): p. 504–510.

40. Jacob, S., et al., The antioxidant α-lipoic acid enhances insulin-stimulated glucose metabolism in insulin-resistant rat skeletal muscle. Diabetes, 1996. 45(8): p. 1024–1029.

41. Lone, Y., et al., DIMETHOATE INDUCED ALTERATION IN SERUM BIOCHEMICAL PARAMETERS IN RATTUS RATTUS. Vol. 6523. 2017. 603–613.

42. Eid, R.A., Apoptosis of rat renal cells by organophosphate pesticide, quinalphos: Ultrastructural study. Saudi Journal of Kidney Diseases and Transplantation, 2017. 28(4): p. 725.

43. Surana, B., J. Mehta, and S. Seshadri, Toxicological effects of QUinolphos and its subsequent reversal by using root extract of Withania somnifera and leaf pulp of Aloe barbadensis. J. Indian Soc. Toxicol, 2009. 4(2): p. 1–5.

44. Hai, D.Q., I. Varga, and B. Matkovics, Effects of an organophosphate on the antioxidant systems of fish tissues. Acta Biologica Hungarica, 1995. 46(1): p. 39–50.

45. Poovala, V.S., H. Huang, and A.K. Salahudeen, Role of reactive oxygen metabolites in organophosphate-bidrin-induced renal tubular cytotoxicity. Journal of the American Society of Nephrology, 1999. 10(8): p. 1746–1752.

46. Takaoka, M., et al., Protective Effect Of α-LIPOIC Acid Against Ischaemic Acute Renal Failure In Rats. Clinical and experimental pharmacology and physiology, 2002. 29(3): p. 189–194.

47. Şehirli, Ö., et al., α-LIPOIC ACID PROTECTS AGAINST RENAL ISCHAEMIA–REPERFUSION INJURY IN RATS. Clinical and experimental pharmacology and physiology, 2008. 35(3): p. 249–255.

48. Koga, H., et al., New α-lipoic acid derivative, DHL-HisZn, ameliorates renal ischemia-reperfusion injury in rats. Journal of Surgical Research, 2012. 174(2): p. 352–358.

49. El-Saad, A.A. and M. Elgerbed, Dimethoate induced hepatotoxicity in rats and the protective roles of vitamin E and N-acetylcysteine. Egypt J. Exp. Biol., 2010. 6(2): p. 219–230.

50. Adham, K., et al., Environmental stress in lake maryut and physiological response of Tilapia zilli Gerv. Journal of Environmental Science & Health Part A, 1997. 32(9-10): p. 2585–2598.

51. Zaahkouk, S.A.M., et al., Carbamate toxicity and protective effect of vit. A and vit. E on some biochemical aspects of male albino rats. Vol. 1. 2000. 60–77.

52. Lucic, A., et al., The effect of dichlorvos treatment on butyrylcholinesterase activity and lipid metabolism in rats. Arhiv za Higijenu Rada I Toksikologiju/Archives of Industrial Hygiene and Toxicology, 2002. 53(4): p. 275–282.

53. Kalender, S., et al., Diazinon-induced hepatotoxicity and protective effect of vitamin E on some biochemical indices and ultrastructural changes. Toxicology, 2005. 211(3): p. 197–206.

54. Kwape, T., et al., Hepato-protective potential of methanol extract of leaf of Ziziphus mucronata (ZMLM) against dimethoate toxicity: biochemical and histological approach. Ghana medical journal, 2013. 47(3): p. 112–120.

55. Morakinyo, A.O., F.O. Awobajo, and O.A. Adegoke, Effects of alpha lipoic acid on blood lipids, renal indices, antioxidant enzymes, insulin and glucose level in streptozotocin-diabetic rats. Biology and Medicine, 2013. 5: p. 26.

56. Zhang, Y., et al., Amelioration of Lipid Abnormalities by α-Lipoic acid Through Antioxidative and Anti-Inflammatory Effects. Obesity, 2011. 19(8): p. 1647–1653.

57. Saltiel, A.R. and C.R. Kahn, Insulin signalling and the regulation of glucose and lipid metabolism. Nature, 2001. 414(6865): p. 799.

58. Budin, S.B., et al., Alpha lipoic acid prevents pancreatic islet cells damage and dyslipidemia in streptozotocin-induced diabetic rats. The Malaysian journal of medical sciences: MJMS, 2007. 14(2): p. 47.

